# AMPK- and cGAS-Mediated Dual Sensing of Glucose and dsDNA Synergizes to Enhance Antitumor Immunity

**DOI:** 10.1101/2025.02.15.638470

**Authors:** Ruyuan Zhou, Jingyan Pei, Shasha Chen, Qin Shen, Qirou Wu, Shengduo Liu, Chen Chen, Xin-Hua Feng, Jian Zou, Tingbo Liang, Y. Jessie Zhang, Sheng-Cai Lin, Penghong Song, Chen-Song Zhang, Qian Zhang, Pinglong Xu

## Abstract

Intratumoral glucose availability is linked considerably to tumor growth, immune evasion, and metastasis. However, the precise molecular mechanisms by which glucose levels govern cellular and immune responses against tumors remain incompletely understood. Here, we explored the context of tumors of intersections between AMP-activated protein kinase (AMPK) and cGAS-STING signaling, a key pathway initiating antitumor immunity. We unexpectedly found that AMPK activation robustly enhances cGAS-STING signaling within cancer and immune cells, which potently impacts melanoma and colorectal cell senescence, organoid apoptosis, and the infiltration of CD4^+^ and CD8^+^ T lymphocytes. Using classic or newly developed AMPK agonist Aldometanib, TBK1^S511E/S511E^ knock-in (KI) mice, glucose metabolic modulator, as well as AMPKα1/α2 dKO melanoma, we demonstrated that dual sensing of intratumoral glucose and dsDNA integrated by the AMPK-TBK1 cascade is essential for initiating antitumor immunity and suppressing cancer progressions. Notably, novel AMPK agonists or intervening in glucose glycolysis by Lonidamine synergized effectively with STING agonists, substantially inhibiting melanoma growth. Therefore, these findings unravel a concise mechanism integrating glucose metabolism and innate DNA sensing into cancer cell fate and propose an effective therapeutic strategy to enhance antitumor immunity.

**HIGHLIGHTS:**

1. Dual and synergistic sensing of glucose and dsDNA facilitates antitumor immunity in melanoma and colorectal cancer;
2. AMPK substitutes intratumoral cGAS-STING-IRF3 signaling through TBK1 phosphorylation at S511;
3. Glucose deficiency induces cancer cell senescence, apoptosis, and T lymphocyte infiltration via the AMPK-TBK1 cascade;
4. Newly developed AMPK agonists synergize with STING agonists to retrieve striking tumor suppression.

## INTRODUCTION

Nucleic acid sensing surveils tissue abnormalities and pathogen invasion and establishes an immune state to restrict microbial infection, modulate adaptive immunity, and guide tissue repair and regeneration (Chan & Gack, 2015, Chen, Sun et al., 2016b, Roers, Hiller et al., 2016, Takeuchi & Akira, 2010, Zhang, Liu et al., 2022b). Pattern recognition receptors (PRRs) such as cGAS (Gao, Ascano et al., 2013, Sun, Wu et al., 2013) monitor microbes and tissue damage by perceiving cytosolic dsDNA derived from pathogens, the mitochondria, or the nucleus (Chen et al., 2016b, Hu & Shu, 2020, Roers et al., 2016, Takeuchi & Akira, 2010). Upon recognition of free dsDNA, cGAS synthesizes 2’3’-cyclic GMP-AMP (cGAMP)(Ablasser, Goldeck et al., 2013, Gao et al., 2013, Wu, Sun et al., 2013, Zhang, Shi et al., 2013), activates STING (also known as MITA or ERIS) (Ishikawa & Barber, 2008, Sun, Li et al., 2009, Zhong, Yang et al., 2008), which triggers the non-canonical STING-PERK pathway in the endoplasmic reticulum (ER) that regulates mRNA translation (Zhang et al., 2022b), induces non-canonical autophagy in the ERGIC (Gui, Yang et al., 2019), and facilitates the activation of TBK1 and IRF3 at the Golgi apparatus (Liu, Cai et al., 2015, Zhang, Shang et al., 2019). Phosphorylated IRF3 translocates into nuclear and transcribes type I interferons (IFN-Is) and numerous IFN-stimulated genes (ISGs) (Fitzgerald, McWhirter et al., 2003, Sharma, tenOever et al., 2003), in coordination with simultaneously activated NF-κB (Chen et al., 2016b, Roers et al., 2016, Takeuchi & Akira, 2010). Therefore, nucleic acid sensing induces mRNA translation arrest (Zhang et al., 2022b), autophagy (Gui et al., 2019, Pilli, Arko-Mensah et al., 2012, Wild, Farhan et al., 2011), phase separation (Meng, Yu et al., 2021), mitochondrial dynamics (Chen, Liu et al., 2020), cell differentiation (Xu, Bailey-Bucktrout et al., 2014), and triggers cell senescence (Dou, Ghosh et al., 2017, Gluck, Guey et al., 2017, Mackenzie, Carroll et al., 2017, Yang, Wang et al., 2017) and cell death (Cerboni, Jeremiah et al., 2017, Melki, Rose et al., 2017, Petrasek, Iracheta-Vellve et al., 2013).

Effects of cGAS-STING signaling in tumor biology are complex and multifaceted (Samson & Ablasser, 2022). cGAS-STING signaling is active in cancerous, immune, and stromal cells, shifting their cellular function, metabolism, and fate. The activation of cGAS-STING signaling releases various IFN-I, cytokines, cGAMP, and DNA, which attracts and matures dendritic cells, macrophages, natural killer cells (NKs), and effector T lymphocytes to mount an antitumor immunity (Chin, Sulpizio et al., 2022, Kwon & Bakhoum, 2020). STING-induced senescence and immune microenvironment modulation also participate in antitumor immunity regulation, positively or negatively ^(Chen & Xu, 2023),(Bakhoum, Ngo et al., 2018),(Chen, Boire et al., 2016a),(Hong, Schubert et al., 2022),(Li, Mirlekar et al., 2022),(Concepcion, Wagner et al., 2022)^, and STING-dependent residency of tumor-associated macrophages (TAMs) attenuate its antitumor role (Zhou, Wang et al., 2024). Therefore, the effects of cGAS-STING in cancer biology could be context- and stage-dependent and should be carefully evaluated during cancer treatments.

Cells in multicellular organisms depend on balanced nutrient availability for survival, growth, and defense. Despite this, the direct molecular connection between nutrient sensing and tissue damage perception remains less understood. AMP-activated protein kinase (AMPK) plays a central role in detecting glucose deficiency and controlling metabolic pathways (Hardie & Ashford, 2014, Hardie, Ross et al., 2012, Lin & Hardie, 2018). AMPK is activated at the lysosomal surface in response to low glucose levels through aldolase and the v-ATPase-Ragulator complex(Zhang, Hawley et al., 2017a, Zhang, Jiang et al., 2014, Zhang, Li et al., 2022a), in addition to being induced by AMP/ATP ratios and kinase such as LKB1 or CaMKII (Garcia & Shaw, 2017, Gonzalez, Hall et al., 2020, Hardie et al., 2012). AMPK masterly regulates an array of cellular processes such as glucose and lipid metabolism, protein synthesis, autophagy, mitochondrial biogenesis, and overall metabolism (Hardie et al., 2012, Mouchiroud, Eichner et al., 2014), and its dysfunction is broadly accompanied by obesity, type 2 diabetes, and cancers. We recently reported that viral infection causes a rapid and dramatic decrease in blood glucose levels and substantially activates AMPK in tissues, which directly phosphorylates TBK1 to trigger IRF3 recruitment and assemble MAVS or STING signalosomes, thus effectively mounting antiviral immunity(Zhang, Liu et al., 2022c). In adipocytes, TBK1 also represses energy expenditure by inhibiting AMPKα (Zhao, Wong et al., 2018). These observations indicate the presence of the intersection between metabolic master kinase AMPK and immune key kinase TBK1. Nevertheless, the roles of metabolism and immunity crosstalk through AMPK-TBK1 signaling in the context of malignant diseases remain poorly understood.

However, hyperglycemia has been considerably linked to diabetes-related vulnerability to malignancy (Klil-Drori, Azoulay et al., 2017), and it impacts cancer immunology in distinct ways. For instance, elevated glucose levels through the hexosamine biosynthesis pathway (HBP) enhance the O-GlcNAcylation level in TAMs, promoting M2-like polarization and cancer progression (Rodrigues Mantuano, Stanczak et al., 2020). In diet-induced obesity, more immunosuppressive myeloid-derived suppressor cells (MDSCs) were observed with higher tumor progression of pancreatic cancer (Clements, Long et al., 2018, Incio, Liu et al., 2016). Meanwhile, tumor cells typically exhibit increased glucose uptake and enhanced glycolysis, which supports key processes for tumorigenesis, such as the biosynthesis of nucleotides, amino acids, and lipids. Focusing on the AMPK-TBK1 axis, here we systematically investigated the link between glucose availability and antitumor immunity. Using the well-documented and newly developed AMPK agonists and TBK1 S511E knock-in (KI) mice, we found that activated AMPK directly phosphorylated intratumoral TBK1 at S511, promoting cGAS-STING-initiated senescence and antitumor immunity, thereby attenuates the progression of melanoma and colorectal cancer. In summary, we proposed an intratumoral dual and synergistic sensing of dsDNA and glucose, which forms a “metabolism and immune sensing” bridge through the concise AMPK-TBK1 signaling and determines tumor progression.

## RESULTS

### Glucose deficiency promotes intratumoral cGAS-STING signaling

To explore the function of glucose metabolism on antitumor immunity in TME, we investigated the state of glucose deficiency and its impacts on cGAS-STING signaling, an innate immune pathway crucial in antitumor immunity. Intratumoral glucose deficiency was evident, as reported (Keerthana, Rayginia et al., 2023, Ramakrishnan, Terry et al., 2025, Reinfeld, Madden et al., 2021), and as indicated by AMPK-induced phosphorylation of metabolic regulator ACC1 in melanoma (Fig. 1A-1B). pACC1 signal was particularly robust at tumor edges, where proliferating tumor cells and immune cells need high glucose demand. cGAMP treatment in primary peritoneal macrophages (PMs) triggered robust activation of TBK1 and IRF3, the central kinase and key signaling mediator of cGAS-STING signaling. Simulating an intratumoral glucose-depleted microenvironment using 2-deoxy-D-glucose (2-DG) resulted in an enhanced cGAS-STING-IRF3 pathway when compared to energy-sufficient conditions, accompanied by robust activation of AMPK (Fig. 1C). We observed a similar enhancement of cGAS-STING signaling in colorectal adenocarcinoma cells (DLD1) and mouse embryonic fibroblasts (MEFs) under energy-deficient conditions (Figs. 1D-1E). AICAR, a well-established AMPK agonist (Corton, Gillespie et al., 1995), expectedly facilitated cGAS-STING signaling in fibroblasts and cancer cells (Figs. 1D-1E). Likewise, glucose depletion amplified mitochondrial DNA (mtDNA)-induced activation of cGAS-STING-IRF3 signaling in colorectal carcinoma cells (Fig. 1F), achieved by combinational treatment of ABT-737, qVD-OPH, and S63845 (AQS) (McArthur, Whitehead et al., 2018, Rongvaux, Jackson et al., 2014, White, McArthur et al., 2014). AICAR also potentiated the sensing of cytosolically exposed dsDNA in DLD-1 cells, simulated by poly(dA:dT) transfection (TpdAdT) (Fig. 1G). These observations across immune cells, fibroblasts, and cancer cells suggest that glucose deficiency, frequently seen in tumors, significantly augments cGAS-STING signaling and is likely through AMPK activation.

**Figure 1.**
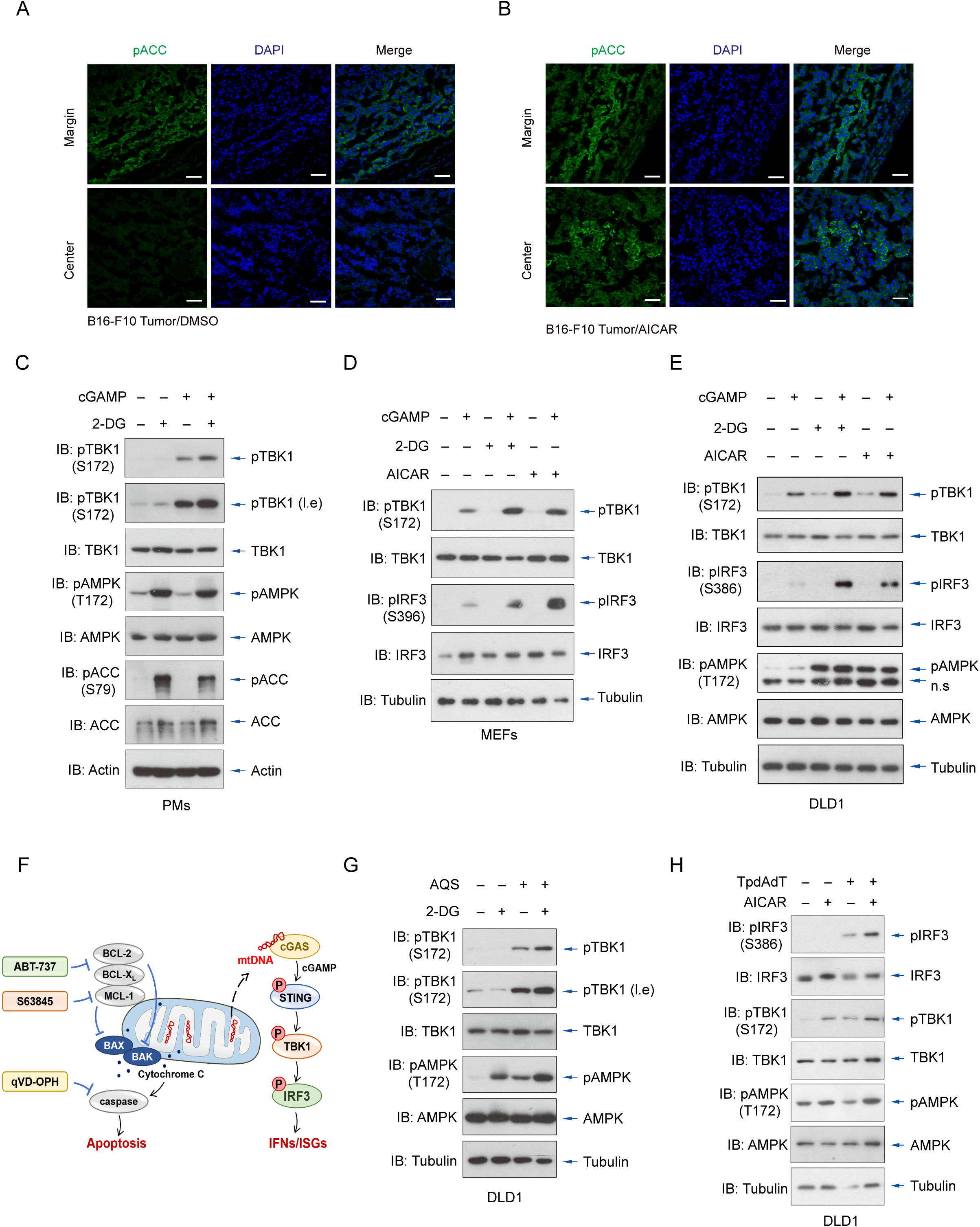
Glucose deficiency promotes intratumoral cGAS-STING signaling. **(A-B)**, The intratumoral glucose deficiency state in B16-F10 melanoma was validated through immunofluorescence of phospho-ACC1 (pACC), a primary substrate of AMPK, by which an evident signal of pACC was seen in the edge of melanoma (A). AMPK agonist AICAR treatment was set as the positive control (B). Scale bars=50 μm**. (C-E),** Cytosolic dsDNA sensing was evaluated in PMs (C), MEFs (D), and DLD1 (E) under 2-DG or AICAR (AMPK agonist) administration in response to STING agonist cGAMP, as indicated by immunoblotting using antibodies recognizing active forms of TBK1 and IRF3. **(F-G)**, Sensing of mtDNAs, released upon the treatment of ABT-737, qVD-OPH, and S63845 (AQS), was similarly augmented by glucose depletion, revealed by the phosphorylation level of TBK1. **(H)**, The effects of AICAR administration on TpdAdT-induced DNA sensing were evaluated in DLD1, representing enhanced dsDNA sensing to double-stranded DNA. Applied to figures 1-6: unless specified, n=3 independent experiments (mean±SEM); *, P<0.05, **, P<0.01, and ***, P<0.001, by statistical analysis of the indicated comparison with ANOVA and Bonferroni correction.

### AMPK facilitates cGAS-STING signaling in cancer cells by TBK1 S511 phosphorylation

To validate that glucose deficiency regulates nucleic acid sensing through AMPK activation, we generated AMPK α1/α2 dKO cells in B16-F10 melanoma cells using a CRISPR-mediated approach (Fig. 2A). Genetic deletion of AMPK catalytic subunits α1/α2 attenuated cGAS-STING-IRF3 signaling in B16 melanoma, as indicated by reduced levels of phospho-STING (S366) and phospho-IRF3 (S396) (Fig. 2A). Likewise, deletion of AMPK α1 and α2 subunits in colorectal carcinoma HCT116 cells similarly decreased RLR-MAVS-IRF3 signaling (Fig. 2B), the dsRNA pathway with an emerged role in antitumor immunity (Chiappinelli, Strissel et al., 2015, Sasaki, Homme et al., 2024). The role of AMPK on TBK1 activation was also evident as AMPK catalytic subunits strongly potentiated the TBK1-IRF3 interaction (Fig. 2C) and TBK1-stimulated IRF3 transactivation (Fig. 2D). Our previous study and the mass spectrometry analyses of AMPK-coexpressed TBK1 have identified that the S511 residue on TBK1 is abundantly modified by AMPKα1 (Zhang et al., 2022c). We found that AMPK inhibitor dorsomorphin (Compound C) abolished TBK1 S511 phosphorylation (Fig. 2E), and an antibody specifically recognizing phospho-TBK1 S511 demonstrated that AMPKα1 directly phosphorylated TBK1 at this residue (Fig. 2F). We thus evaluated the capability of wild-type, phospho-resistant (S511A), and phosphomimetic TBK1 (S511E) to activate IRF3 by an IRF3-responsive reporter, which indicated that AMPK-mediated TBK1 phosphorylation at S511 was critical for driving IRF3 transactivation (Fig. 2G). Expectedly, a compromised interaction between phospho-resistant TBK1 and IRF3 was seen (Fig. 2H). These findings suggest that AMPK enhances cGAS-STING signaling by phosphorylating TBK1 at S511 to prime TBK1 activation.

**Figure 2.**
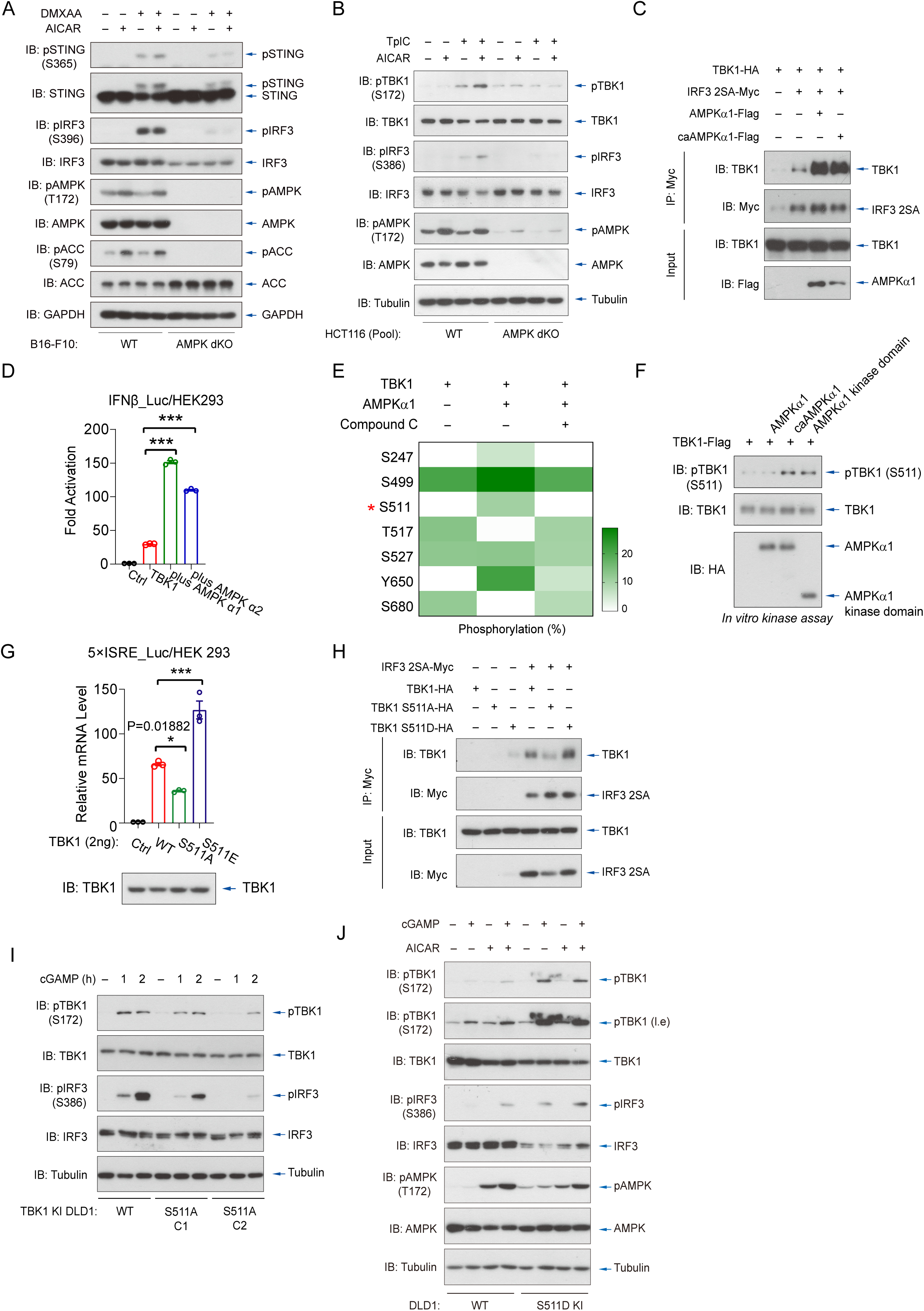
AMPK facilitates cGAS-STING signaling in cancer cells by TBK1 S511 phosphorylation. **(A)**, Double knockout (dKO) of AMPK α1 and α2 impaired the activation of TBK1 and IRF3 following DMXAA stimulation in B16-F10 melanoma cells. **(B)**, The genetic ablation of AMPK α1 and α2 subunits reduced TpIC-induced cytosolic RNA sensing in HCT116, revealed by a significant decrease in phosphorylation of TBK1 and IRF3. **(C)**, Coimmunoprecipitation assay revealed coexpression of AMPKα1 or caAMPKα1 (activated AMPKα1) enhanced this association of TBK1 and IRF3 2SA. **(D)**, Reporter assay showed that the expression of AMPK α1 and α2 subunits strongly promoted TBK1-stimulated IRF3 transactivation with IFNβ promoter in HEK293 cells. **(E)**, Mass spectrometry analysis identified that AMPKα1 phosphorylated the S511 residue of TBK1, and the phosphorylation was completely inhibited by the AMPK inhibitor dorsomorphin (Compound C). **(F)**, caAMPKα1 and AMPKa1 kinase domain directly phosphorylated TBK1 at S511 in vitro kinase assay. **(G-H)**, TBK1 mutants S511A (resistant to AMPK modification) exhibited significantly reduced activity IRF3-responsive reporter assays (G) and a weaker interaction with IRF3 2SA (H). Meanwhile, TBK1 S511E (mimics AMPK-phosphorylated TBK1) demonstrated a strong potential to bind and activate IRF3. **(I-J)**, Generated DLD1 knock-in (KI) clones with mutations that either prevent (S511A) or mimic (S511D) AMPK-mediated modification using CRISPR-based genome editing. cGAMP-stimulated cGAS-STING pathway was weakened in TBK1 S511A KI cells (I) but strengthened in S511D KI cells (J).

To validate this AMPK-TBK1 axis in cancer cells, we generated and examined knock-in (KI) clones of colorectal cells DLD1 with mutations that either prevent (S511A) or mimic (S511D) AMPK-mediated modification by CRISPR-based genome editing (Zhang et al., 2022c). cGAS-STING signaling was attenuated when endogenous TBK1 was mutated into the S511A variant that resists AMPK modification (Fig. 2I). Conversely, STING signaling was robustly activated when endogenous TBK1 was mutated to the S511D variant that mimics AMPK-TBK1 phosphorylation, to a level comparable to those under AICAR treatment (Fig. 2J). Noticeably, the AICAR-enhanced activation of STING signaling was absent in these DLD1 KI cells (Fig. 2J). These observations suggest that AMPK-mediated TBK1 phosphorylation at residue S511 is critical for cancer cells to respond to glucose deficiency.

### AMPK drives STING-induced cancer cell senescence and apoptosis

cGAS-STING responds to dsDNA leakage and drives cellular senescence, which could be important in tumor growth arrest and potentially in antitumor immunity (Dou et al., 2017, Gluck et al., 2017, Yang et al., 2017). We found that upon chemical induction of DNA damage, colorectal carcinoma DLD1 cells displayed strong phenotypes of cellular senescence, evidenced by SA-β-Gal staining and senescence-associated secretory phenotypes (SASPs) (Fig. 3A-3B). Genetic ablation of AMPK α1/α2 subunits endowed DLD1 with apparent resistance to DNA damage-induced cellular senescence (Fig. 3A). By contrast, AICAR facilitated the progression of these colorectal cancer cells into senescence (Fig. 3B). In agreement with these observations, simulating the glucose-depleted microenvironment using 2-DG greatly enhanced DLD1 senescence (Fig. 3C). We further found that by utilizing a zebrafish model of damage-induced senescence, AMPK activation by AICAR promoted camptothecin (CPT)-induced senescence in zebrafish, indicated by increased numbers of SA-β-Gal-positive cells in the heads and tails of zebrafish (Fig. 3D). Inhibition of AMPK by dorsomorphin strongly attenuated CPT-induced senescence, as expected (Fig. 3D). These observations suggest that AMPK is critical to regulate damage-induced cellular senescence, mediated by cGAS-STING signaling.

**Figure 3.**
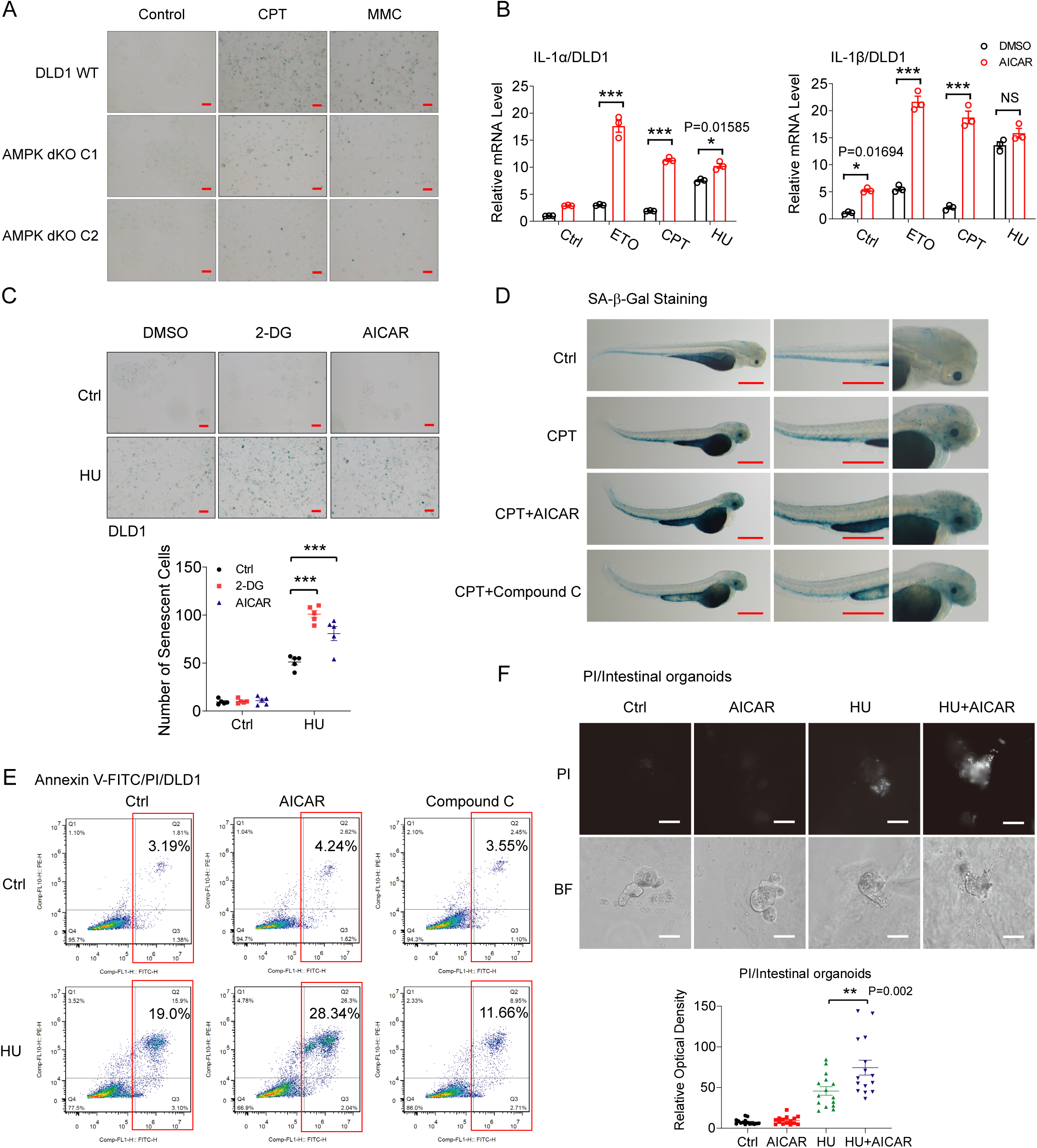
AMPK drives STING-induced senescence and apoptosis of cancer cells and organoids. **(A**-**B)**, Chemical induction of DNA damage in DLD1 gut epithelial cells by CPT, mitomycin C (MMC), etoposide (ETO), and hydroxyurea (HU) treatment led to a robust phenotype of cellular senescence, as assessed by SA-β-Gal staining (A) and SASP features, including the mRNA expression of IL1α and IL1β (B). Genetic ablation of AMPKα1/α2 in DLD1 cells prevented DNA sensing-initiated cellular senescence (A). Conversely, AICAR treatment significantly enhanced DNA damage-induced cellular senescence (B). Scale bars = 100 μm. **(C)**, Energy deficiency stimulation of AMPK by 2-DG or AICAR led to an exaggerated phenotype of cellular senescence in DLD1 cells, particularly upon HU-induced DNA damage. Scale bars = 100 μm. ***, P<0.001, by ANOVA with Bonferroni correction. **(D**, ) Cellular senescence was assessed in zebrafish embryos (n=30) subjected to chemical-induced damage by incubation with CPT (50 nM) and to SA-β-Gal staining. A strong phenotype of cellular senescence was detected in the heads and tails of zebrafish; this phenotype was mildly upregulated by AICAR treatment but was mitigated by dorsomorphin. Scale bars = 500 μm. **(E)**, FACS analyses of early and late apoptotic cells among DLD1 cells revealed that activation of AMPK by AICAR promoted cell apoptosis induced by cytosolic DNA sensing, which triggered by DNA damage inducer HU, but inhibition of AMPK by dorsomorphin suppressed HU-induced apoptosis. **(F)**, Intestinal crypts were isolated from mice at 6 weeks of age and cultured in vitro in the absence or presence of the AMPK agonist AICAR. PI staining revealed that cytosolic DNA sensing stimulated by the DNA damage inducer HU resulted in robust apoptosis of intestinal cells in organoids, and AICAR exaggerated this effect. Scale bars=50 μm. n=15 intestinal organoids examined (mean±SEM). *, P=0.002, compared with wild-type organoids (ANOVA with Bonferroni correction).

The activation of cytosolic dsDNA sensing also mediates a context-dependent apoptotic program (Gulen, Koch et al., 2017), an evolved strategy beneficial to tissue homeostasis and immune modulation. We found that activation of cGAS-STING by DNA damage inducer HU (Wu, Zhang et al., 2019) triggered significant levels of colorectal cancer cell apoptosis, which was noticeably regulated by AICAR and dorsomorphin (Fig. 3E). Additionally, by utilizing in vitro-cultured intestinal organoids, we detected a strong phenotype of HU-induced apoptosis among intestinal cells, which was further exaggerated by AMPK activation (Fig. 3F). These observations suggest that glucose deficiency and resulting AMPK activation controls both senescence and apoptosis of cancer and functional cells through an AMPK-TBK1 axis.

### AMPK-TBK1 signaling facilitates STING-mediated antitumor immunity in melanomas

Next, we utilized a naturally occurring and constitutively active STING SAVI mutant (R281Q) to generate B16-F10 melanoma with inducible and constitutively active STING signaling. Upon implanting subcutaneously into wild-type C57BL/6 mice, we found that melanomas harboring STING R281Q SAVI mutation exhibited significantly reduced growth (Figs. 4A-4C), attributed mainly from STING-induced expression of type I interferons (IFNs) (Wu et al., 2019). Increased immune cells infiltrated these melanomas with constitutive STING signaling, particularly T cells (Figs. 4D-4E), indicating enhanced antitumor immunity. Administration of AMPK agonist AICAR significantly intensified T lymphocyte infiltration in melanomas (Figs. 4D-4E) and suppressed tumor growth substantially (Figs. 4A-4C), suggesting that AMPK can amplify STING-mediated antitumor immunity in melanoma.

**Figure 4.**
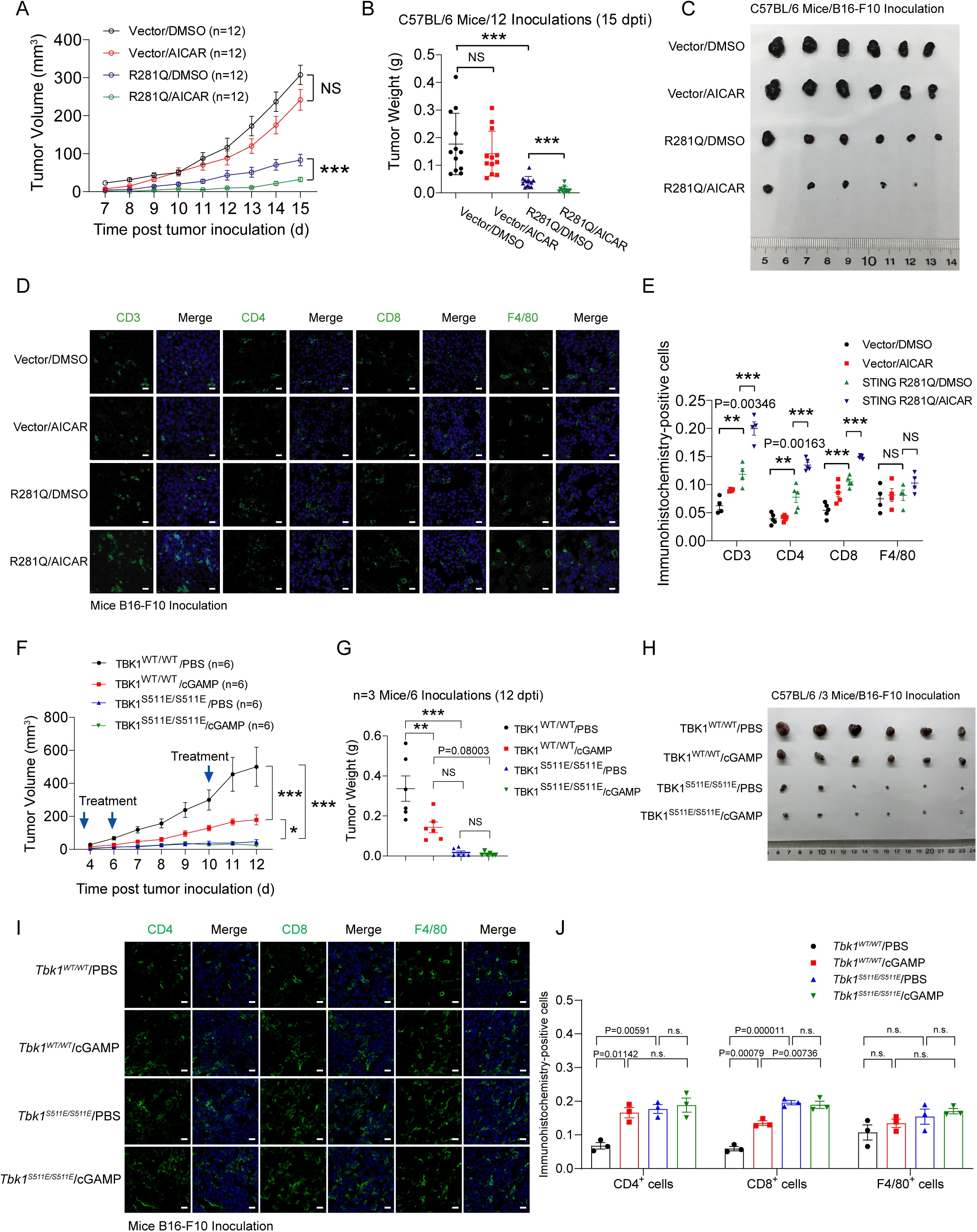
AMPK-TBK1 signaling facilitates STING-mediated antitumor immunity in melanomas. **(A-C)**, The STING SAVI mutant (R281Q) was constitutively active (caSTING). The volumes (A), weights (B), and photos (C) of the subcutaneously B16-F10 melanoma growth in wild-type C57BL/6 mice revealed a markedly decreased growth in melanomas expressing the STING SAVI mutant (R281Q). AICAR administration further increased this effect of STING signaling, resulting in robust tumor growth arrest. n=12 inoculations per group (mean ± SEM), measured 7 days post tumor inoculation (dpti). **(D-E)**, The administration of AICAR with caSTING profoundly increased immune cell infiltration, particularly the infiltration of the CD4^+^ and CD8^+^ T cells into melanomas, as detected by Immunohistochemistry (IHC) upon STING SAVI mutant expression (D). The statistics for the immunohistochemistry-positive cells were calculated (E). Scale bars=20 μm. **, P<0.01, and ***, P<0.001, by ANOVA with Bonferroni correction. **(F-H)**, B16-F10 melanoma cells were implanted subcutaneously into TBK1^WT/WT^ or TBK1^S511E/S511E^ C57BL/6 mice with intratumoral injection of cGAMP. Volumes (F), weights (G), and photos (H) of melanoma revealed that melanoma growth was markedly attenuated in these TBK1^S511E/S511E^ mice. **(I-J)**, Immunohistochemistry (I), and statistics (J) showed B16-F10 melanoma from TBK1^S511E/S511E^ mice accompanied with the increased immune cell infiltration. Stimulation in TBK1^S511E/S511E^ mice by cGAMP did not further enhance the infiltrated T cells and antitumor immunity effect. Scale bars=20 μm.

To determine whether the AMPK-TBK1 axis mediates this enhanced antitumor immunity effects, we generated knock-in (KI) mice harboring the TBK1 S511E mutation using a CRISPR-mediated strategy (Zhang et al., 2022c), which mimics AMPK-mediated TBK1 phosphorylation. The TBK1 S511E homozygous mice were viable, fertile, and appeared phenotypically normal. However, melanoma growth was markedly attenuated in these TBK1 S511E mice (Figs. 4F-4H), along with a significantly increased level of T lymphocyte infiltration (Figs. 4I-4J). More elevated antitumor immunity was not seen in TBK1 S511E knock-in mice in the presence of cGAMP (Figs. 4F-4H), further suggesting the importance of AMPK-TBK1 signaling in this process. These consistent observations propose the AMPK-TBK1 axis as a critical factor in facilitating STING-mediated antitumor immunity.

### AMPK agonists synergize well with cGAMP to promote antitumor immunity in melanomas

To evaluate the potential for improving intratumoral antitumor immunity through this AMPK-TBK1 axis, we employed in the in vivo model of melanoma a well-characterized AMPK agonist AICAR ^(Corton^ ^et^ ^al., 1995)^ and a newly developed AMPK agonist Aldometanib ^(Zhang^ ^et^ ^al., 2022a)^, which acts through distinct mechanisms. As expected, administration of cGAMP in mice, a natural STING ligand, resulted in significant tumor growth suppression. AICAR (Figs. 5A-5C) and Aldometanib (Figs. 5D-F) markedly enhanced melanoma growth arrest, synergizing well with cGAMP. These findings suggest that maneuvering AMPK activation can effectively strengthen STING-induced antitumor immunity. Intriguingly, we observed that Aldometanib also enhanced antitumor effects in mice bearing AMPK dKO B16-F10 melanomas (Figs. 5G-5I), suggesting an additional cancer cell-nonautonomous effect of Aldometanib in melanoma.

**Figure 5.**
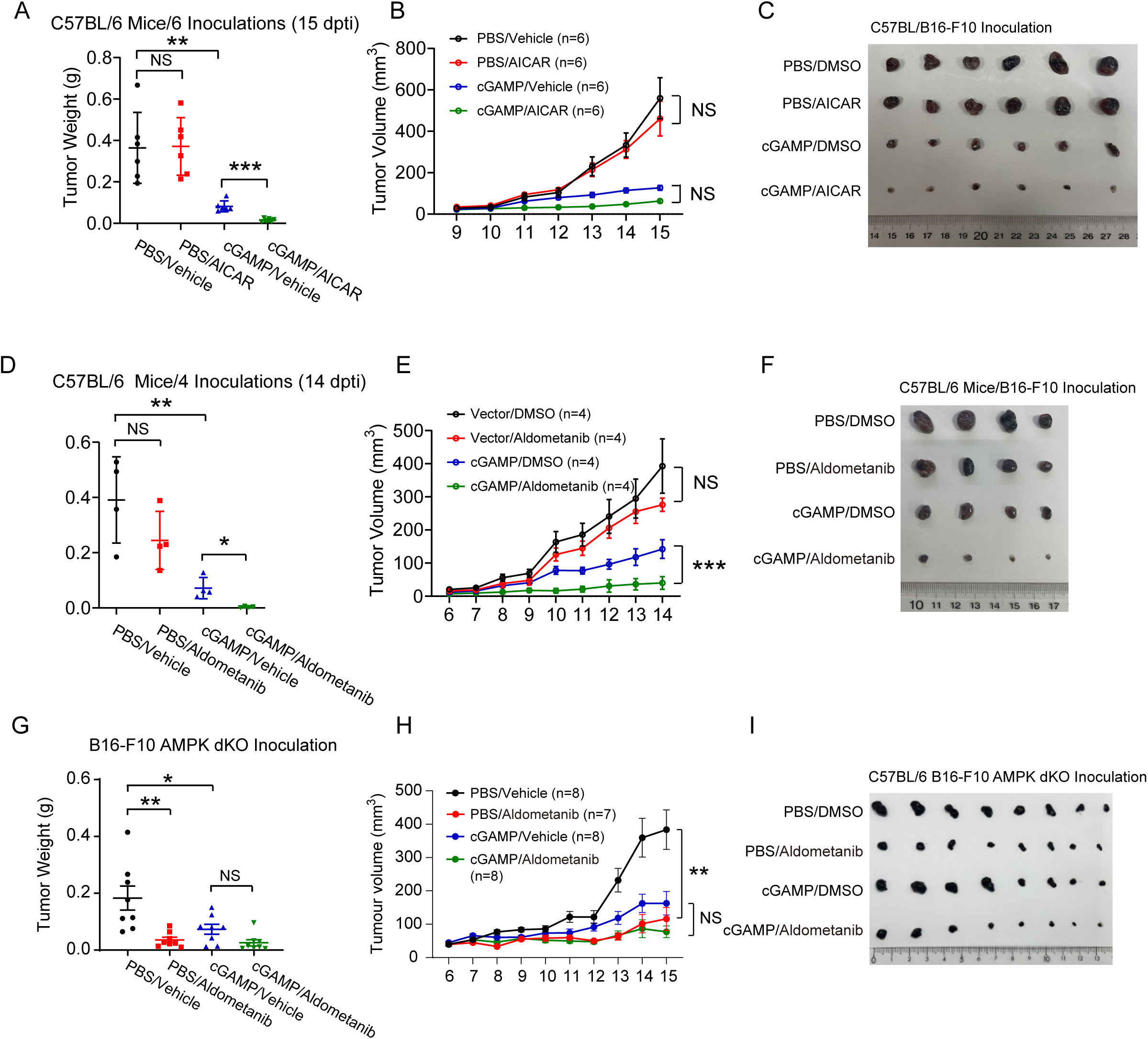
AMPK agonists synergize well with cGAMP to promote antitumor immunity in melanomas. **(A-C)**, The weights (A), volumes (B), and photos (C) of the subcutaneously B16-F10 melanoma growth in wild-type C57BL/6 mice with administration of cGAMP or AICAR. AICAR administration further promoted the antitumor immunity effect of cGAMP-induced STING signaling, inhibiting tumor growth. n=6 inoculations per group (mean ± SEM). **(D-F)**, The weights (D), volumes (E), and photos (F) of the subcutaneously B16-F10 melanoma growth in wild-type C57BL/6 mice with administration of Aldometanib and cGAMP. Adding AICAR enhanced the suppression of tumor growth induced by cGAMP. n=4 inoculations per group (mean ± SEM). **(G-I)**, Administration of Aldometanib alone inhibited the growth of AMPK dKO B16-F10 tumors in WT mice. The weights (G), volumes (H), and photos (I) showed similar tumor sizes of AMPK dKO B16-F10 tumors with the treatment of Aldometanib or cGAMP. n=8 inoculations per group (mean ± SEM).

### Interrupting aerobic glycolysis potentiates antitumor immunity through the AMPK-TBK1 cascade

Next, we interrupted glucose metabolism by inhibiting aerobic glycolysis in cancer cells using Lonidamine, a classical hexokinase inhibitor (Floridi, Paggi et al., 1981), to evaluate the potential of glucose metabolism intervention in antitumor immunity. Lonidamine interferes with energy metabolism and induces AMPK activation (Roberts, Tan-Sah et al., 2014, Wu, Zhang et al., 2023, Zhang et al., 2017a). Notably, we found that combinational use of cGAMP and Lonidamine resulted in substantial tumor growth arrest (Figs. 6A-6C), along with significantly increased infiltration of intratumoral T lymphocytes in melanomas (Figs. 6D-6E). Furthermore, we found that Lonidamine promoted the robust activation of intratumoral cGAMP-induced cGAS-STING signaling in B16-F10 melanomas, evidenced clearly by increased levels of STING clustering, phosphorylated STING (pSTING S365), and phosphorylated TBK1 (pTBK1 S172) (Fig. 6F). These findings suggest that targeting glucose metabolism enhances STING-mediated immunity and can be a promising strategy in antitumor therapeutic, such as in melanomas.

**Figure 6.**
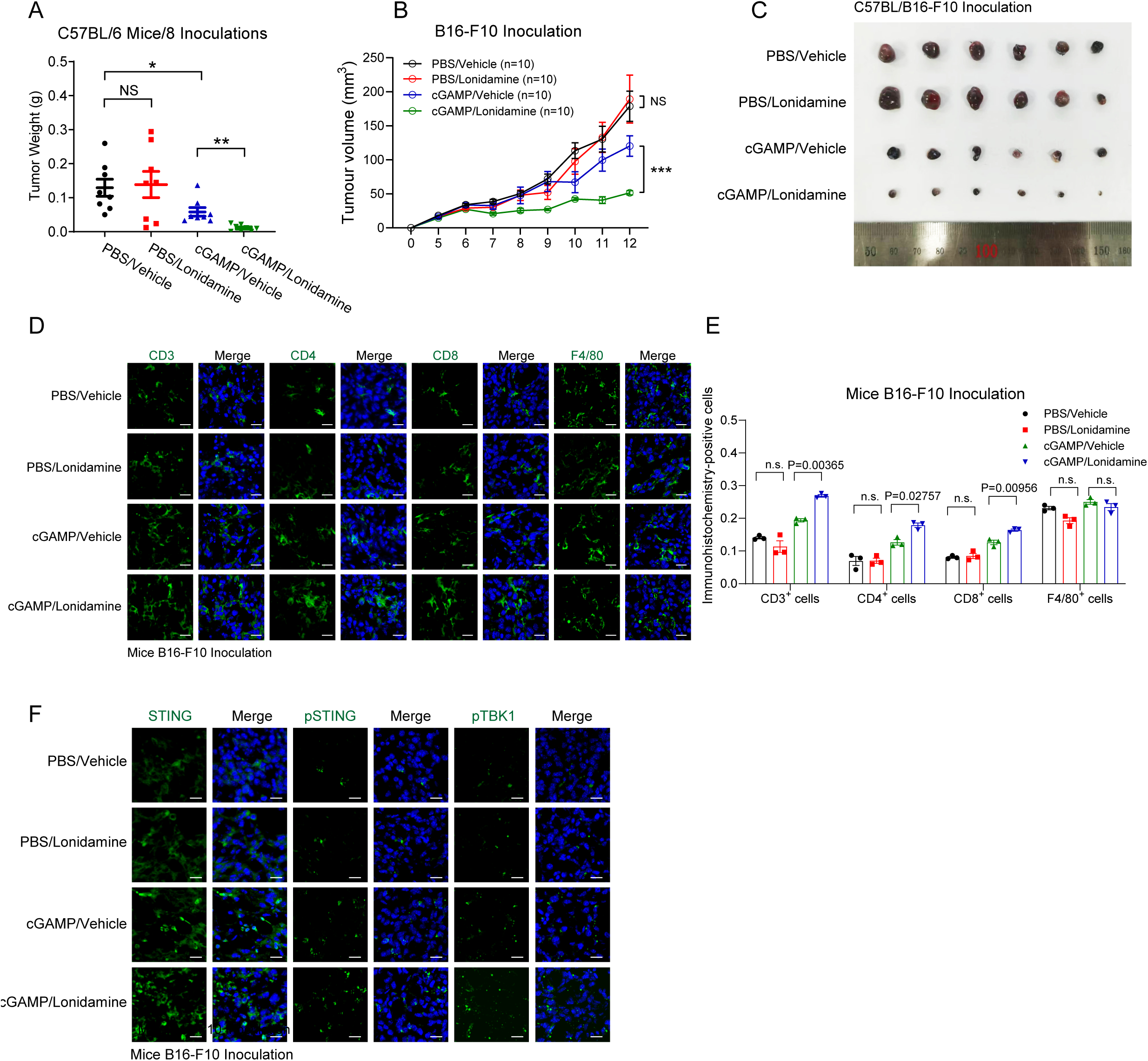
Disrupting aerobic glycolysis potentiates antitumor immunity through the AMPK-TBK1 cascade. **(A-C)**, The administration of Lonidamine (Hexokinase inhibitor) with cGAMP profoundly inhibited the growth of B16-F10 melanoma, revealed by the weights (A), volume curve (B), and images (C) of B16-F10 melanoma. n=8 or 10 inoculations per group (mean ± SEM). **(D-E)**, Enhanced infiltration of CD3^+^, CD4^+,^ and CD8^+^ T lymphocytes were imaged in B16-F10 melanomas with the administration of Lonidamine and cGAMP; Scale bars, 20 μm. (H) n=3 per group (mean ± SEM). **(I)**, Immunohistochemistry showed that Lonidamine treatment enhanced cGAMP-induced cGAS-STING signaling in B16-F10 melanomas, as evidenced by increased levels of phosphorylated STING (pSTING S365) and phosphorylated TBK1 (pTBK1 S172), and total intratumoral STING protein levels; Scale bars=20 μm.

## DISCUSSION

Tumor cell metabolism depletes essential nutrients in the tumor microenvironment (TME), leading to significant glucose competition between cancer and immune cells. Meanwhile, the intervention of cGAS-STING signaling is a significant feature of cancers, mainly by genetic deletion of signaling components and suppression of signaling cascades, which attenuates its beneficial roles in antitumor immunity (Ghosh, Saha et al., 2021, Meng et al., 2021, Wu et al., 2019). However, the exact mechanisms for cGAS-STING activation and its contribution to cellular behaviors in cancer and immune cells, such as cell death, senescence, IFN secretion, and metabolic changes, need further interrogation. This study elucidated how the glucose sensor AMPK senses intratumoral glucose abundance and promotes antitumor immunity through direct phosphorylation and activation of TBK1, an essential kinase during innate immune responses. Given the complex energy metabolism in TME, understanding the AMPK-TBK1 axis opens new avenues for targeting glucose metabolism in both cancer and immune cells. Through activating cGAS-STING signaling and enhancing T-cell-mediated antitumor immunity, these findings provide significant therapeutic targets for drug development in cancer immunotherapy.

Cancer cells primarily rely on aerobic glycolysis for energy metabolism, through which glucose is converted into lactic acid. This metabolic shift exacerbates glucose deficiency stress within the tumor, leading to metabolic competition between tumor cells and immune cells. Notably, these metabolic changes in immune cells contribute to establishing an immunosuppressive microenvironment for accelerating tumor growth and immune evasion. While previous studies have extensively characterized the Warburg effect and its impact on tumor proliferation and immune suppression, this study focuses on how metabolic sensing mechanisms combined with nucleic acid sensing influence the immune landscape of the TME. The cGAS-STING pathway facilitates immune cell recruitment and inhibits tumor progression through cellular senescence and apoptosis (Chen & Xu, 2023). A recent study has highlighted that the AMPK-TBK1 axis plays a crucial role in innate antiviral immunity, extending AMPK’s function beyond its classical roles in glucose and lipid metabolism, protein synthesis, autophagy, and mitochondrial function (Zhang et al., 2022c). Changes in glucose metabolism in the TME are closely linked to immune responses. We found that the interaction between AMPK and cGAS-STING forms a “metabolic and immune sensing” bridge, which synergistically senses the metabolic state of tumor cells and tumor microenvironment to decide the magnitude of immune responses during tumorigenesis. AMPK directly phosphorylates S511 residue on TBK1, enhancing STING-IRF3 signaling activation and promoting T-cell infiltration into the TME, significantly inhibiting melanoma growth. Therefore, these findings expand the understanding of metabolic integration in innate immunity and provide new insights into antitumor immunity.

New therapeutics for tumor treatment increasingly focus on regulating cellular metabolism and immune responses. We propose that modulating glucose levels within tumors, the activation state of AMPK, and the modulation of AMPK-TBK1 signaling could offer novel aspects for antitumor therapeutics. As a critical energy sensor, AMPK is activated in response to glucose deprivation, thereby adjusting the cellular metabolic state. Our study revealed that AMPK activation leads to direct phosphorylation of TBK1 at S511, enhancing cGAS-STING-IRF3 signaling and promoting antitumor immune responses and STING-induced cellular senescence and apoptosis, thereby potently inhibiting tumor growth. Besides, AMPK plays an important role in the adaptive stress response of tumor cells, regulating key metabolic pathways to inhibit tumor cell proliferation (Lin & Hardie, 2018). We supposed AMPK also influences the metabolism of immune cells in the TME, particularly macrophage polarization, which impacts tumor immune evasion. By modulating AMPK activity, immune cell surveillance of tumors can be enhanced through cGAS-STING signaling, effectively preventing tumor immune escape.

In conclusion, our study reveals the key role of the concise AMPK-TBK1 axis in regulating intratumoral glucose metabolism and promoting antitumor immune responses, which considerably connects glucose metabolism with innate immunity. We propose that targeting the AMPK-TBK1 axis, the intersection between intratumoral glucose metabolism and antitumor immunity, could reach a substantial synergy for cancer treatments.

## ACKNOWLEDGMENTS

This research was sponsored by the NSFC Projects (32321002, 32430028, and 31830052 to P.X., 82271768 to Q.Z., and 82001668, 32370759 to S.C.) and the National Key Research and Development Program of China (2021YFA1301401 to P.X. and 2021YFD1801103 to Q.Z.). Thanks also to technical assistance by the Life Sciences Institute core facilities, Zhejiang University.

## AUTHOR CONTRIBUTIONS

Q.Z. and R.Z. carried out most experiments. C.-S.Z., J.P., S.C., Q.S., Q.W., S.L., and C.C. contributed to several experiments. X.-H.F., J.Z., T.L., Y.J.Z., S.-C.L., P.S., and C.-S.Z. helped with reagents, data analyses, and discussions. P.X., Q.Z., and R.Z. conceived the study and experimental design and wrote the manuscript.

## COMPETING FINANCIAL INTERESTS

The authors declare no competing financial interests.

## MATERIALS AND METHODS

### Mice

TBK1 S511E transgenic mice were on C57BL/6 background, and both male and female littermates were used in all the experiments. GemPharmatech generated TBK1 Knock-in transgenic C57BL/6 mice. To create point mutations (S511E) at mouse Tbk1 locus by CRISPR/Cas9-mediated genome engineering, Tbk1 S511E (TCT to GAA) specific sgRNA oligos were designed by the CRISPR website (http://crispr.mit.edu/) targeting sequence at S511 locus (5’-TTCTATTGTTCCCTGAGAAC -3’) and (5’-ATTTAGCTTTCCAGTTCTCA -3’), with S511E oligo donor sequence (5’-CATTTGGATCTGATCCGTTGTTCTGACCTAACCTAACCCGTTGTATTTAGCTTTCCAGCGAACAGGGAACAATAGAAAGCAGTCTTCAGGACATCAGCAGCAGGCTGTCTCCAGGGGGCT -3’). Cas9 mRNA and sgRNA generated by in vitro transcription and donor oligo were co-injected into fertilized eggs for knock-in mouse production. The target region of the mouse Tbk1 locus was sequenced to confirm targeting. All the animals were bred and maintained in a pathogen-free animal facility at the laboratory animal center of Zhejiang University. The care of experimental animals was approved by the Zhejiang University committee and followed Zhejiang University guidelines.

### Zebrafish

Zebrafish AB wild-type embryos (male/female) were raised at 28.5°C in E3 egg water. The care of experimental animals was approved by the Zhejiang University committee and followed Zhejiang University guidelines.

### Peritoneal Macrophages

Peritoneal macrophages were isolated from C57BL/6 mice at 6-8 weeks of age with the Brewer thioglycollate medium (Sigma-Aldrich)-induced approach. Three days after the intraperitoneal injection of 2.5 mL of 3% thioglycollate medium, peritoneal macrophages were isolated and cultured in the RPMI 1640 medium.

### Cell Lines

HEK293, HCT116, DLD1, MEFs, and B16-F10 cells were from ATCC. No cell lines used in this study were found in the database of commonly misidentified cell lines maintained by ICLAC and NCBI Biosample. Cell lines were frequently checked in morphology under microscopy and tested for mycoplasma contamination but were not authenticated. All cell lines were cultured in DMEM medium with 10% fetal bovine serum (FBS) at 37°C in 5% CO_2_ (v/v), except for peritoneal macrophages that were maintained in RPMI 1640 medium,

### Intestinal Organoids

Isolated small intestines were opened longitudinally, washed three times with cold PBS, removed the villi by scraping, and incubated twice in cold PBS with 21mM EDTA for 151min on ice. After removing the EDTA medium, the fractions were incubated in 50 mL cold PBS with a fierce shake and passed through a 70-μm cell strainer (BD Bioscience) to remove the residual villous material. Isolated crypts were centrifuged at 300 g for 5 min to separate the crypts with single cells. The centrifuged fractions were suspended and cultured in Matrigel with medium (Advanced DMEM/F12 supplemented with penicillin/streptomycin, 10 mM HEPES, Glutamax, 13N2, 13B27 (all from Invitrogen), and 1mM N-acetylcysteine (Sigma) containing growth factors 50 ng mL^-1^ EGF, 100 ng mL^-1^ noggin, 1mg mL^-1^ R-spondin). The organoid formation was observed and analyzed every day after plating.

## METHOD DETAILS

### Expression Plasmids, Viruses, Reagents, and Antibodies

Expression plasmids encoding Flag-, Myc-, or HA-tagged wild-type or mutations of human TBK1, IRF3, and the reporters of IFNβ_Luc and 5xISRE_Luc have been described previously (Liu, Chen et al., 2017, Zhang, Meng et al., 2017b). Flag- or HA-tagged human AMPKα1, AMPKα2, caAMPKα1 and AMPKα1 kinase domains were constructed using the pRK5 mammalian expression vector. Site-directed mutagenesis about TBK1 S511A/D/E and AMPKα1 T172D were generated by PCR-based cloning performed by a kit from Stratagene. All coding sequences were verified by DNA sequencing, and the detailed information for construction was provided in the attached Supplementary Table 2 and upon requirement.

The pharmacological reagent, Compound C (Selleck), AICAR (Selleck), 991 (Selleck), A-769662 (Selleck), ABT-737 (Selleck), qVD-OPH (Selleck), S63845 (Selleck), MK2206 (Selleck), Doxycycline (Sangon Biotech), 2-DG (Sangon Biotech), cGAMP (Invivogen), poly(I:C) (Invivogen), poly (dA:dT) (Invivogen), puromycin (Yeasen), G418 (Yeasen), Hydroxyurea (HU) (Sigma), and Lonidamine (Selleck) were purchased and used according to manual instructions. Aldometanib was generated by Dr. Sheng-Cai Lin (Xiamen University, Xiamen).

The monoclonal anti-TBK1 (3504S, 1:5,000 dilution), anti-pTBK1(S172) (5483S, 1:3,000 dilution), anti-IRF3 (4302S, 1:2,000 dilution), anti-pIRF3 (S396) (4947S, 1:5,000 dilution), anti-AMPKα (5831S, 1:1,000 dilution), anti-pAMPKα (T172) (50081S, 1:1,000 dilution), anti-pACC (S79) (3661S, 1:1,000 dilution), anti-ACC (3676S, 1:1,000 dilution), anti-Myc (2276S, 1:3,000 dilution), and anti-HA (3724S, 1:5,000 dilution) antibodies were purchased from Cell Signaling Technology. The anti-pIRF3 (S386) (ab76493, 1:3,000 dilution), anti-STING (ab181125, 1:100 dilution) were purchased from Abcam, and the anti-α-tubulin (T6199, 1:10,000 dilution), anti-β-actin (A5441, 1:10,000 dilution), anti-GAPDH (EPR16891, 1:3000), anti-HA (H9658, 1:200 dilution), and anti-Flag (M2) (F3165, 1:5,000 dilution) were purchased from Sigma. The anti-rabbit IgG and anti-mouse IgG antibodies were purchased from Santa Cruz. The anti-pTBK1 (S511) was generated in collaboration with Abcam, targeting the phospho-TBK1 S511.

### Plasmids Transfection and Virus Infection of Cultured Cells

Lipofectamine 3000 (Invitrogen), Polyethyleneimine (PEI, Polysciences), or Lipofectamine RNAiMAX (Invitrogen) transfection reagents were used for the transfection of plasmids, poly(dA:dT), and poly(I:C). Digitonin (Sigma) was used to induce cGAMP.

### CRISPR/Cas9-mediated Generation of AMPK**α**1/**α**2**^-/-^** and TBK1 Knock-in Cells

Guide RNA sequences targeting AMPKα1 and α2 mRNA sequence (hAMPKα1, 5’-AAAGTTTGAGTGCTCAGAAG -3’, 5’-GTGATGGAATATGTCTCAGG -3’; hAMPKα2, 5’-TCAATTAACAGGCCATAAAG -3’, 5’-GTTATTTAAGAAGATCCGAG -3’), TBK1 S511A/D knock-in mRNA sequence (hTBK1, 5’-TATTTAGCTTTCCAGTTCTC -3’, 5’-TTCTATTGTTCCCTGAGAAC -3’) were used to clone the genes into the vector pX330, which were transfected into DLD1 cells by LipofectAmine 3000, together with the pCas9-2A-GFP plasmid, or with pCas9-2A-GFP and pMD18 in the case of TBK1 S511A/D knock-in generation. 36 hours after transfection, cells with green fluorescence were sorted with a Flow Cytometer (BD FACS Aria II) and propagated. Immunoblotting with anti-AMPK(or the sequencing of the genomic PCR products) identified clones. All gRNAs used in the experiments were also attached in Supplementary Table 1.

### Luciferase Reporter Assay

HEK293 cells were transfected with indicated reporters (100 ng) bearing an open read frame (ORF) coding Firefly luciferase, along with the pRL-Luc with Renilla luciferase coding as the internal control for transfection and other expression vectors specified in the results section. In brief, after 24 hours post-transfection, cells were lysed by passive lysis buffer (Promega), and luciferase assays were performed using a dual luciferase assay kit (Promega), quantified with POLARstar Omega (BMG Labtech) and normalized to the internal Renilla luciferase control.

### Quantitative RT-PCR Assay

The DLD1 cells stimulated with Hydroxyurea (HU), Camptothecin (CPT), or Etoposide (ETO) were lysed, and total RNA was extracted using an RNAeasy extraction kit (Axygen). cDNA was generated by a one-step iScript cDNA synthesis kit (Vazyme), and quantitative real-time PCR was performed using the EvaGreen Qpcr MasterMix (Abm) and CFX96 real-time PCR system (Bio-Rad). Relative quantification was expressed as 2-ΔCt, where Ct is the difference between the primary Ct value of triplicates of the sample and an endogenous L19 mRNA control. The primer sequences used are listed in the following:

hIL-1α, 5’-AGTGCTGCTGAAGGAGATGCCTGA -3’, 5’-CCCCTGCCAAGCACACCCAGTA -3’;

hIL-1β, 5’-ATGATGGCTTATTACAGTGGCAA -3’, 5’-GTCGGAGATTCGTAGCTGGA -3’;

RPL19, 5’-ATGTATCACAGCCTGTACCTG -3’, 5’-TTCTTGGTCTCTTCCTCCTTG -3’;

All primers used in the qRT-PCR assay were also attached in Supplementary Table 1.

### Coimmunoprecipitations and Immunoblottings

HEK293 transfected with specified plasmids encoding Myc-, Flag-, or HA-tagged AMPKα1, AMPKα2, TBK1s, and IRF3s was lysed using the modified MLB lysis buffer (20 mM Tris-Cl, 200 mM NaCl, 10 mM NaF, 1 mM NaV_2_O_4_, 1% NP-40, 20 mM β-glycerophosphate, and protease inhibitor, pH 7.5) (Xu et al., 2014). Cell lysates were then subjected to immunoprecipitation using the antibodies of anti-Flag (Sigma, F3165-5MG, 1:200 dilution), anti-Myc (CST, 2276S, 1:200 dilution), or anti-HA (Sigma, H9658, 1:200 dilution) for transfected or induced proteins. After 3-4 washes with the MLB, adsorbed proteins were resolved by SDS-PAGE (Bio-Rad) and immunoblotting with the indicated antibodies. Cell lysates were also analyzed using SDS-PAGE and immunoblotting to control the protein abundance.

### *In vitro* Kinase Assay

HEK293 cells were transfected with plasmids encoding Flag-, Myc-, or HA-tagged TBK1s or AMPKα and lysed by the modified MLB lysis buffer after 24 hours of transfection. Immunoprecipitations were performed by using anti-Flag (Sigma, F3165-5MG, 1:200 dilution), anti-Myc (2276S, 1:200 dilution), or anti-HA (CST, 3724S, 1:200 dilution) antibodies. With two washes by the MLB and two washes by the kinase assay buffer (200 μM ATP, 200 μM AMP, 20 mM Tris-HCl, 1 mM EGTA, 5 mM MgCl_2_, 0.02% 2-mercapto-Ethanol, 0.03% Brij-35, and 0.2 mg mL^-1^ BSA, PH 7.4), immunoprecipitated TBK1s and AMPKα1 were incubated in the kinase assay buffer at 30°C for 60 min on THERMO-SHAKER (1400 rpm/min). The reaction was stopped by adding 2 × SDS loading buffer and subjected to SDS-PAGE and specified immunoblotting.

### Cellular senescence assays

To evaluate damage-induced cellular senescence, cells at approximately 60-70% confluence were treated with Hydroxyurea (HU, 10 mM), Camptothecin (CPT, 1 µM), Etoposide (ETO, 25 µM) or Mitomycin (MMC, 1 µM) for 24 hours, replaced with fresh medium, and cultured for another 5 days and harvested for SA-β-Gal assay, by using a cellular senescence assay kit (Yeasen 40754ES60) and according to the manufacturer’s manual. Images were acquired using a converted fluorescence microscope from Nikon., cells were lysed after 5-8 days stimulations and subjected to RNA extraction and qRT-PCR assays as described in the previous section, to detect the senescence-associated secretory phenotype (SASPs), such as the expression of cytokines including IL-1α and IL-1β.

### Senescence assays in zebrafish

Zebrafish embryos were treated with 50 nM CPT with/without 1mM AICAR or 10 µM Compound C from 1.5 days post fertilization (dpf) to 3.5 dpf and then fixed in 4% paraformaldehyde in PBS at 4°C. After 3 washes with PBS (pH 7.4 and pH 6.0) at 4°C, SA-β-Gal staining was performed according to the manual of the senescence assay kit (Yeasen 40754ES60). Animals were photographed under a dissecting microscope.

### Murine allograft growth of B16 melanoma

C57BL/6 wild-type mice were maintained under specific-pathogen-free (SPF) conditions and randomly selected for tumor injection. Six- to eight-week-old mice were administered a subcutaneous injection of 1.5×10^6^ B16-F10 melanoma cells and treated with or without AICAR (5 mg/kg) or once a day, Aldometanib was supplied either through oral gavaging(Zhang et al., 2022a). From the ninth-day post-tumor inoculation, the injected mice were monitored for tumor growth daily as described in the protocols approved by the IACUC of Zhejiang University. Other treatment regimens included: cGAMP, 2⍰µg intratumoral injection every 3 days, starting on day 5 post-implantation; Lonidamine, 25 mg/kg intraperitoneal injection every 3 days, starting on day 4 post-implantation. Tumor size is presented as a square caliper measurement and was calculated based on two perpendicular diameters (mm^2^). The maximum diameter of tumors is limited up to 10 mm by ethical permission, upon which the mice were sacrificed and considered as dead due to tumor burden.

### Murine tumor Immunohistochemistry

For immunohistochemistry assays, tumor samples were dissected, fixed in 4% paraformaldehyde for 12 hours at 4°C, dehydrated with 30% sucrose overnight at 4°C, embedded in optimal cutting temperature compound, and immediately frozen at -80°C. The sectioned samples at 10 μm thickness were washed two times with PBS, permeabilized with 0.5% Triton X-100, blocked in 3% horse serum in PBS for 30 min, and incubated sequentially with primary antibodies, including anti-CD3 (Abcam, 16669, 1:100 dilution), anti-CD4 (eBioscience, 14976680, 1:100 dilution), anti-CD8 (eBioscience, 14080880, 1:100 dilution), and anti-F4/80 (Bio-Rad, MCA497RT, 1:100 dilution). After the sections were incubated with Alexa-labeled secondary antibodies (Jackson, 111-095-003, 1:500 dilutions, Invitrogen, A11006, 1:1000 dilutions) and extensively washed, they were stained with DAPI (Santa Cruz Biotech) and mounted with ProLong^TM^ Gold antifade reagent (Invitrogen). Immunofluorescence images were obtained and analyzed using a Zeiss LSM710 or LSM 880 confocal microscope.

### Nano-liquid Chromatography/Tandem MS (Nano LC-MS/MS) Analysis

Phoenix National Proteomics Core services performed Nano LC/tandem MS analysis for protein identification and characterization and label-free quantification. Tryptic peptides were separated on a C18 column and analyzed by LTQ-Orbitrap Velos (Thermo). Proteins were identified using the National Center for Biotechnology Information search engine against the human or mouse RefSeq protein databases. The mass spectrometry proteomics data of TBK1 modifications by AMPK have been deposited to the ProteomeXchange Consortium (http://proteomecentral.proteomexchange.org) via the iProX partner repository with the dataset identifier PXD033926.

## QUANTIFICATION AND STATISTICAL ANALYSIS

Quantitative data are presented as the mean ± standard error of the mean (SEM) from at least three independent experiments. When appropriate, the statistically significant differences between multiple comparisons were analyzed using the one-way or two-way ANOVA test with Bonferroni correction. Differences were considered significant at p<0.05. All samples were included in the analyses if preserved and properly processed, and no samples or animals were excluded except for zebrafish with conventional injection damage. No statistical method was used to predetermine sample size, and all experiments except those involving animals were not randomized. Immunoblotting, reporter assays, and qRT-PCR experiments have been repeated a minimum of three times independently to ensure reproducibility. The investigators were not blinded to allocation during experiments and outcome assessment.

---

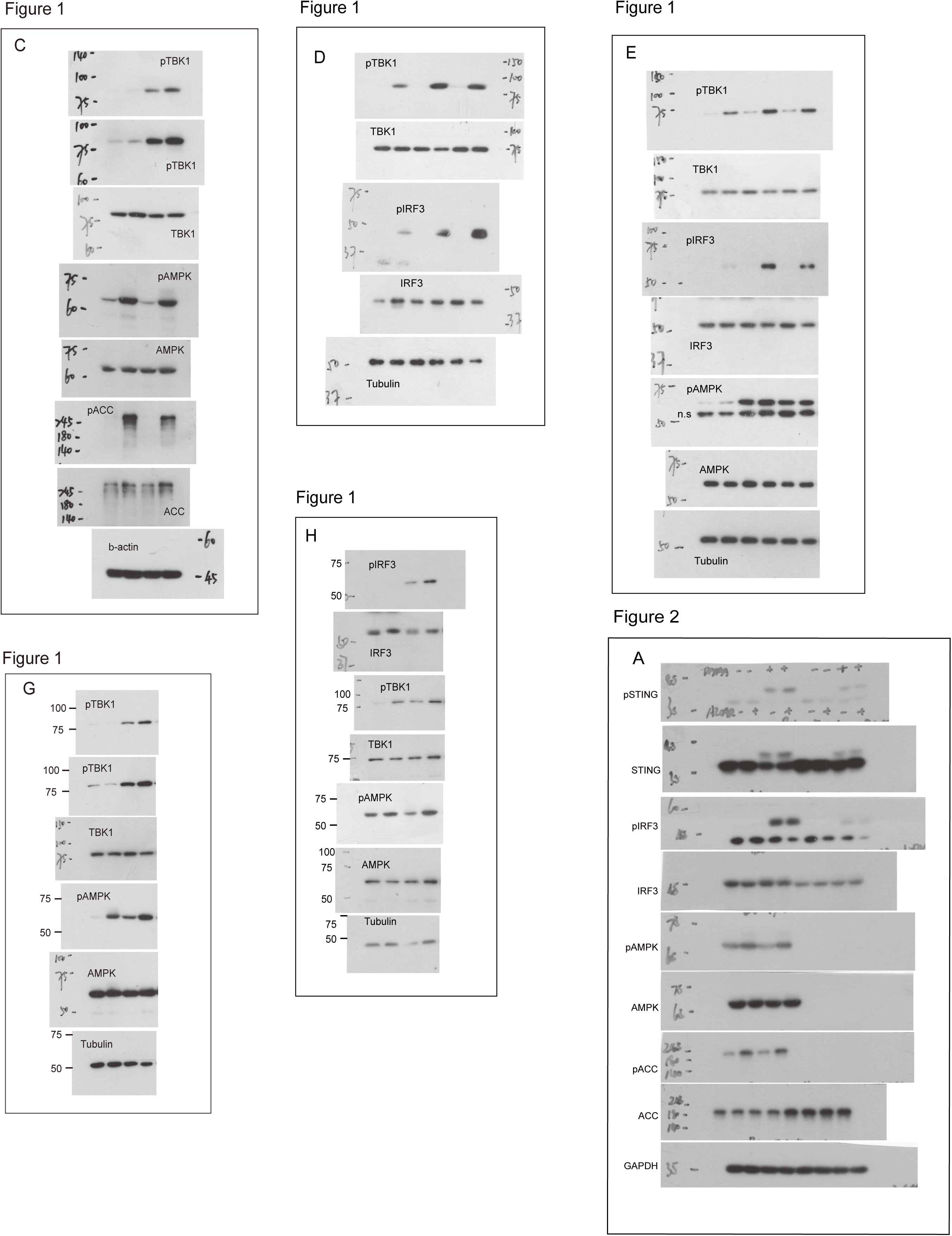

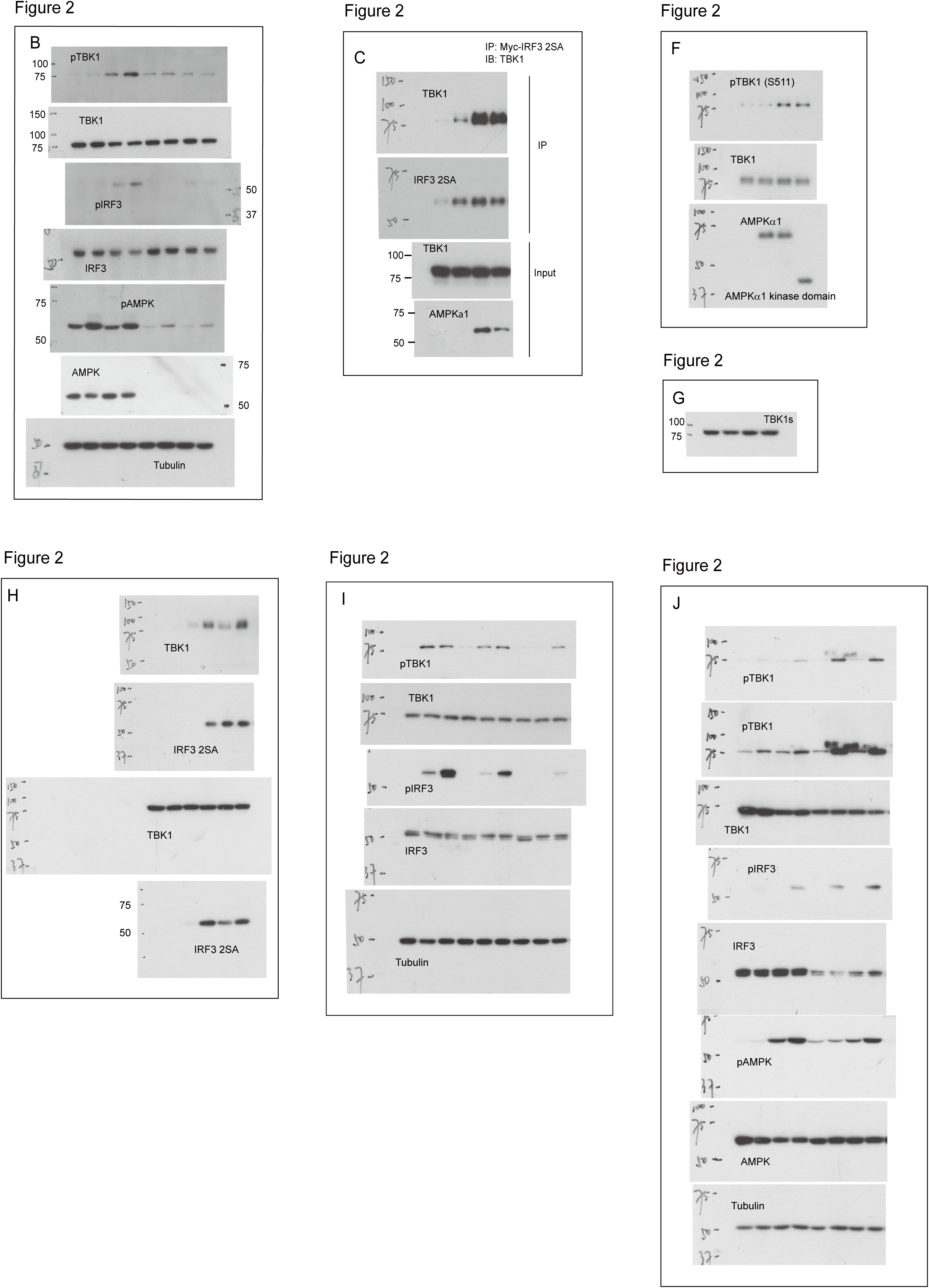

**Suppl. Table 1.**
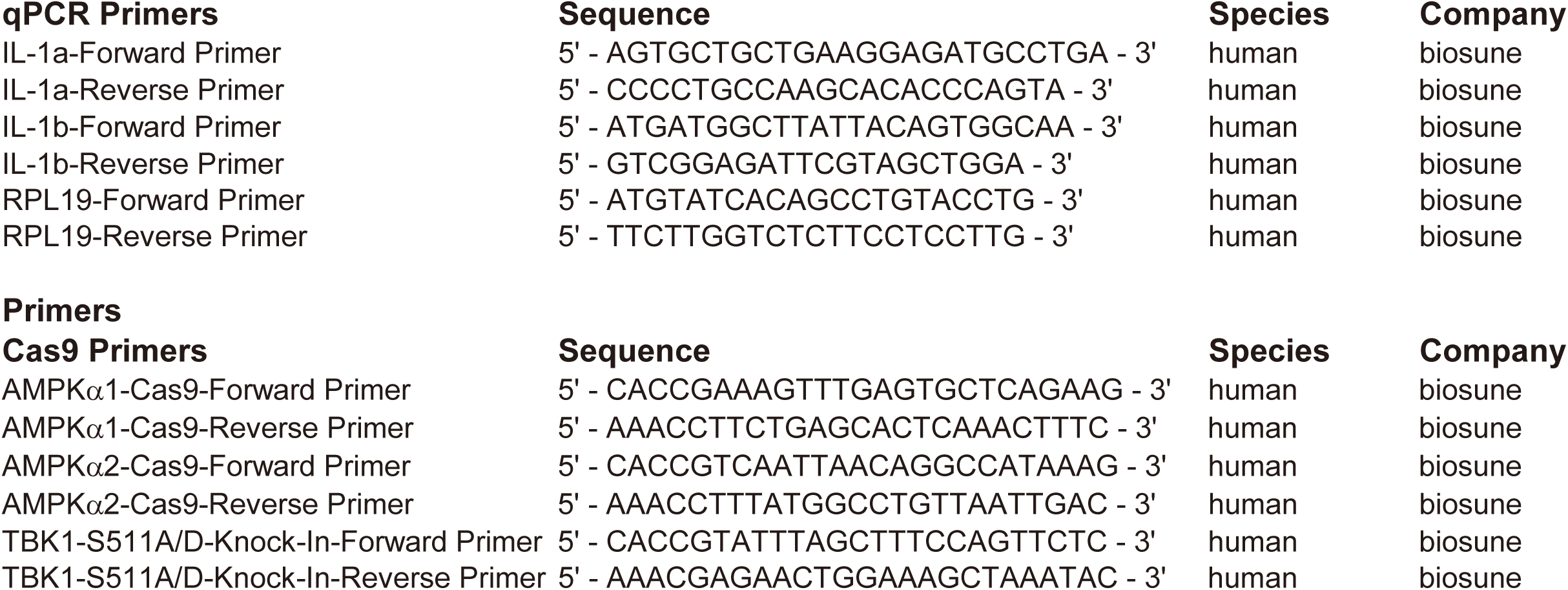
Oligos Used in Study.

**Suppl. Table 2.**
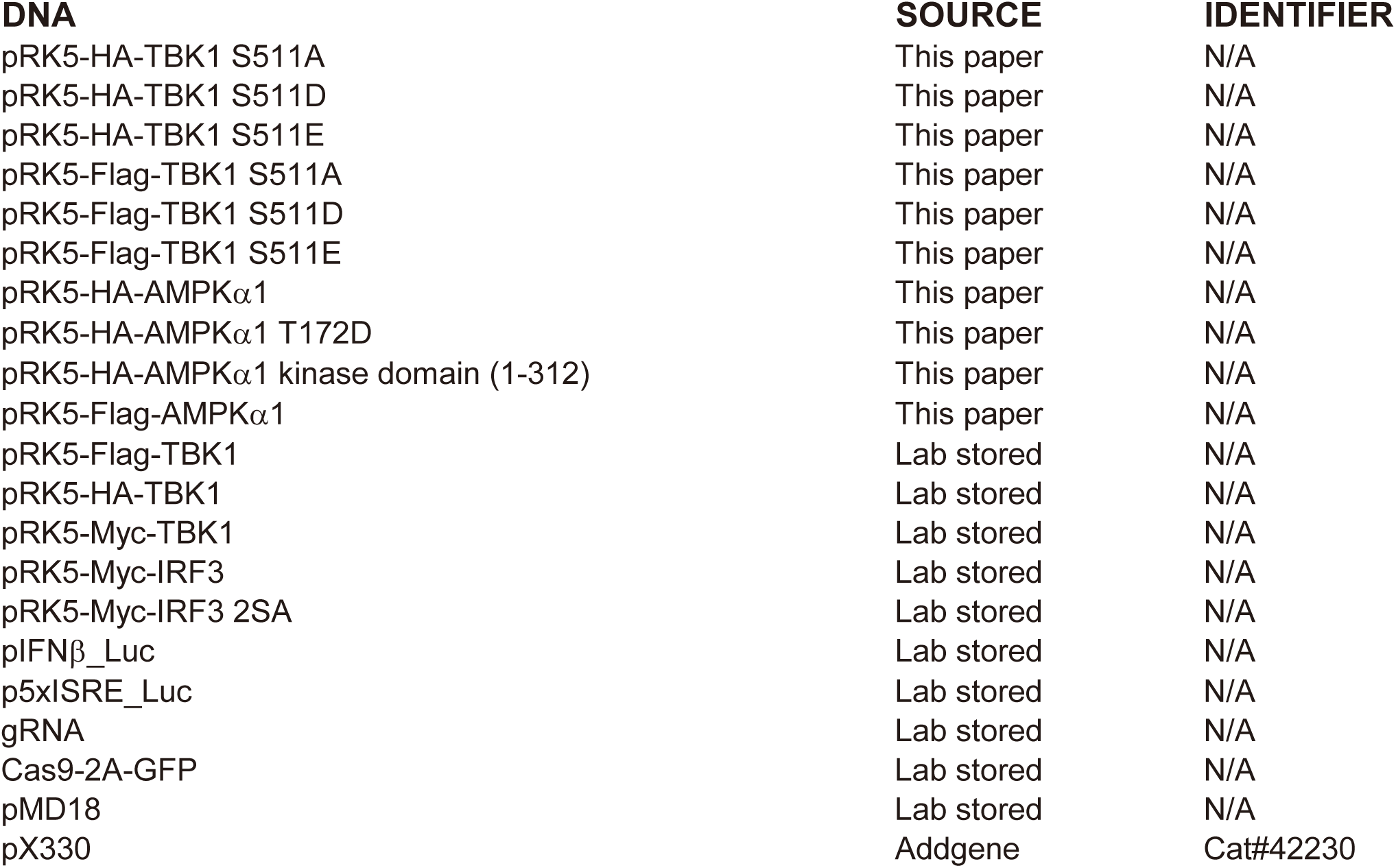
List of Recombinant DNA.

| **DNA** | **SOURCE** | **IDENTIFIER** |
| --- | --- | --- |
| pRK5-HA-TBK1 S511A | This paper | N/A |
| pRK5-HA-TBK1 S511D | This paper | N/A |
| pRK5-HA-TBK1 S511E | This paper | N/A |
| pRK5-Flag-TBK1 S511A | This paper | N/A |
| pRK5-Flag-TBK1 S511D | This paper | N/A |
| pRK5-Flag-TBK1 S511E | This paper | N/A |
| pRK5-HA-AMPKα1 | This paper | N/A |
| pRK5-HA-AMPKα1 T172D | This paper | N/A |
| pRK5-HA-AMPKα1 kinase domain (1-312) | This paper | N/A |
| pRK5-Flag-AMPKα1 | This paper | N/A |
| pRK5-Flag-TBK1 | Lab stored | N/A |
| pRK5-HA-TBK1 | Lab stored | N/A |
| pRK5-Myc-TBK1 | Lab stored | N/A |
| pRK5-Myc-IRF3 | Lab stored | N/A |
| pRK5-Myc-IRF3 2SA | Lab stored | N/A |
| pIFNβ_Luc | Lab stored | N/A |
| p5xISRE_Luc | Lab stored | N/A |
| gRNA | Lab stored | N/A |
| Cas9-2A-GFP | Lab stored | N/A |
| pMD18 | Lab stored | N/A |
| pX330 | Addgene | Cat#42230 |

**Suppl. Table 3.** Statistics Source Data. (See details in the following sheets)

## Notes

### Competing Interest Statement

The authors have declared no competing interest.

### Summary of Updates

Corrections of author list and grammar errors.

